# Pneumatic Nano-Sieve for CRISPR-based Detection of Drug-resistant Bacteria

**DOI:** 10.1101/2023.08.17.553737

**Authors:** Ruonan Peng, Xinye Chen, Fengjun Xu, Richard Hailstone, Yujie Men, Ke Du

## Abstract

The increasing prevalence of antibiotic-resistant bacterial infections, particularly methicillin-resistant Staphylococcus aureus (MRSA), presents a significant public health concern. Timely detection of MRSA is crucial to enable prompt medical intervention, limit its spread, and reduce antimicrobial resistance. Here, we introduce a miniaturized nano-sieve device featuring a pneumatically-regulated chamber for highly efficient MRSA purification from human plasma samples. By using packed magnetic beads as a filter and leveraging the deformability of the nano-sieve channel, we achieve an on-chip concentration factor of 15 for MRSA. We integrated this device with recombinase polymerase amplification (RPA) and clustered regularly interspaced short palindromic repeats (CRISPR)-Cas detection system, resulting in an on-chip limit of detection (LOD) of approximately 100 CFU/mL. This developed approach provides a rapid, precise, and centrifuge-free solution suitable for point-of-care diagnostics, with the potential to significantly improve patient outcomes in resource-limited medical conditions.

## Introduction

The prevalence of antibiotic-resistant bacterial infections has become a major concern for both individuals and healthcare facilities, causing an estimated 1.27 million fatalities globally and contributing to nearly 5 million deaths in 2019 ^1^. One particular pathogen, methicillin-resistant *Staphylococcus aureus* (MRSA), stands out as a prominent multidrug-resistant (MDR) bacterium that presents a serious challenge ^2^. MRSA can cause skin infections ^3^, pneumonia ^4^, and even sepsis ^5^, and it exhibits resistance to beta-lactam antibiotics, including methicillin, penicillin, amoxicillin, and oxacillin, which are commonly used in the treatment of bacterial infections ^6^. Consequently, infections caused by MRSA are associated with high morbidity and mortality rates, making it a significant public health issue ^7^.

Early detection of MRSA is crucial as it enables timely and appropriate medical intervention, prevents the spread of these pathogens, and reduces the risk of antimicrobial resistance ^8^. For detection, it is essential to isolate bacteria from the samples collected from nose ^9^, blood ^10^ or urine ^11^. Membrane-based filtration is a widely used and advantageous method for capturing bacteria due to its cost-effectiveness, simplicity, and rapidness ^12^. However, when dealing with blood samples, the presence of blood cells would cause membrane clogging issues ^13,14^. Furthermore, a significant challenge associated with this technique is the extensive volume of iterative washing buffer required to retrieve the captured bacteria from the membrane ^15^. This washing process can inadvertently lead to dilution of the captured bacteria, reducing their concentration to levels that may fall below the detection limit of downstream detection processes. Alternatively, microfluidic platforms have emerged as a powerful tool in the field of bacterial purification and concentration ^16,17^. Those platforms could be functionalized to rapidly and efficiently separate and concentrate target bacteria, depending on various working principles, including inertial force ^18,19^, hydrodynamics ^20,21^, electrophoresis ^22,23^, and acoustics ^24,25^. However, these techniques require either complicated fabrication processes or extra laboratory-based instruments, increasing the complexity of microfluidic platforms for delicate operations. Therefore, it is important to develop a simple and direct process of fabricating a microfluidic platform while ensuring it can effectively separate the target bacteria. Traditional MRSA detection methods, such as cultured-based techniques, are time-consuming and labor-intensive. Molecular methods like polymerase chain reaction (PCR) require thermocyclers and sophisticated bulky equipment, which renders them unsuitable for resource-limited point-of-care (POC) environments ^26^. Clustered regularly interspaced short palindromic repeats (CRISPR)-Cas (CRISPR-associated) systems, particularly the Cas12 and Cas13 nucleases, have gained significant attention in the field of in vitro diagnostics ^27^. Among them, Cas12a relies solely on a complementary crRNA for targeting specific DNA sequences and utilizes a single RuvC domain to cut the target DNA, a process known as cis-cleavage ^28,29^. Moreover, Cas12a exhibits collateral activity, referred to as trans-cleavage, which allows it to non-specifically cleave neighboring single-stranded DNAs (ssDNA) following target binding ^30^. To exploit this feature for detection purposes, ssDNA can be labeled with a fluorophore-quencher, and upon Cas12a activation through target binding, the cleavage of ssDNA generates an increase in fluorescence signal ^31^. CRISPR-Cas12a system operates at 37°C, making it more suitable for POC detection than traditional PCR methods ^32^. Additionally, by combining the CRISPR-Cas system with isothermal amplification methods such as recombinase polymerase amplification (RPA) ^33^, rolling circle amplification (RCA) ^34^, and loop-mediated isothermal amplification (LAMP) ^35^, the specificity and sensitivity of CRISPR-Cas detection can be further enhanced in POC settings ^36^. Notably, RPA stands out as a widely adopted amplification method due to its simplicity, rapidity, and compatibility with the same temperature requirements as CRISPR assays.

Herein, we introduce a miniaturized and versatile nano-sieve device with a pneumatically-regulated chamber that allows for rapid purification and highly concentrated isolation of MRSA from plasma samples. To achieve this, we developed a simplified, direct, and cost-efficient fabrication process for this nano-sieve device, incorporating multiple channels. With a three dimensional (3-D) magnetic beads-stacked microstructure within the channel, the highly efficient bacteria capture was preceded by precisely controlling the applied flow rate. Leveraging the deformability of the nano-sieve channel, a remarkable concentration (around 15-fold) of captured bacteria was achieved by adjusting the volume ratio of initial sample solution and retrieved buffer solution. This unique functionality of nano-sieve significantly enhances the limit of detection (LOD) when combined with the developed RPA and CRISPR-Cas assay, ultimately achieving an on-chip LOD of approximately 100 CFU/mL. Importantly, the entire process can be completed in less than 4 hours under physiological temperature and room temperature, without the need of centrifugation. Therefore, our approach of integrating the microfluidic-based multiplexing purification and a rapid and precise molecular detection could potentially improve the sensitivity and specificity of MRSA detection.

## Results and Discussion

The schematic of the whole system is presented in **Fig. 1a**, where multiple nano-sieve channels are designed for multiplexing separation of bacteria from initial samples. **Fig. 1b** shows the picture of a practical nano-sieve device, including the beads-stacked channel filled with blue food dye and the pneumatic layer filled with red food dye. This indicates the device can be successfully fabricated without any leaking issues. In **Fig. 1c**, the fabrication flow of a pneumatically-regulated nano-sieve device is exhibited. It started with a thin layer of TEOS (200 nm in thickness) deposited onto a pre-cleaned glass wafer, which was followed by a spin-coated layer of positive photoresist. After that, a pattern of nano-sieve channel was transferred from a plastic photomask to the photoresist layer by photolithography technique, then defined on the layer of TEOS by BOE process. The patterned channel with a thickness of 200 nm was created, and finally covered by a thin layer of positive photoresist as a sacrificial support for PDMS bonding procedure. This coated photoresist can eliminate the technical issue of collapsed PDMS roof ^37^, significantly enhancing the fabrication of nano-fluidic channels. The pneumatic chamber was made by employing a 3-D printed mold, and a thin film of uncured PDMS was sandwiched by glass slides, with a supporting ORACAL film to define the thickness of this PDMS film to be created. Then the pneumatic chamber and the cured PDMS thin film were bonded through the plasma treatment. The fabrication of the nano-sieve device was subsequently completed by bonding the pneumatic chamber layer and the glass substrate patterned with nano-sieve channels via plasma treatment. Both treatments (marked by the red dashed rectangles) were followed by the baking process on a hot plate to achieve a strong bonding in between.

**Fig. 1.**
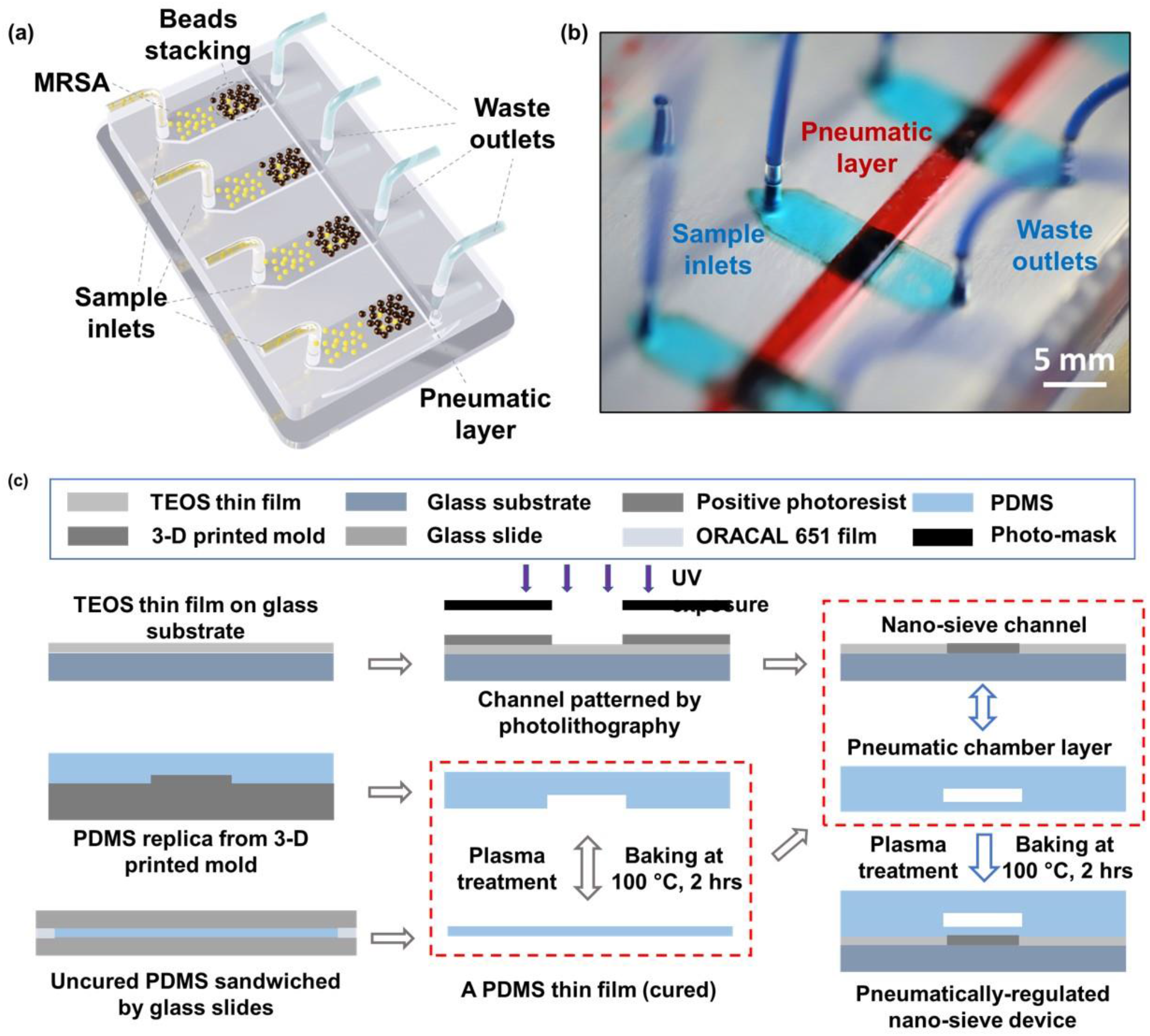
(a) Schematic of nano-sieve device. The MRSA sample is injected through sample inlets, and subsequently, bacteria become entrapped within the beads stacking region of the microfluidic channel, allowing for effective MRSA concentration. Meanwhile, the waste liquid, consisting of plasma and phosphate-buffered saline (PBS), exits the device through designated waste outlets. (b) Image of realized multi-channel nano-sieve with food dye. (c) Fabrication process of nano-sieve device.

The optical micrograph in **Fig. 2a** displays the pre-loaded stacked beads array within the half section of the nano-sieve channel, which are well secured by the positive pressure applied in the pneumatic chamber. Another half section per channel was connected to the outlets for collecting the waste liquids. **Fig. 2b** presents the experimental setup, regarding a multiplexing separation of target bacteria under the observation of fluorescence microscopy. Within these nano-sieve channels as shown in **Fig. 2c**, only the target bacteria stained by the green dye can display the green fluorescent signals. During the flow condition, the bacteria were carried by flowing fluid, moving forward to the area of stacked beads, where they were physically captured by the array of 5 μm beads. As shown in **Fig. 2d** and **Fig. 2e**, the original bacterial sample and retrieved bacterial sample were compared to highlight the on-chip concentration capability of this powerful nano-sieve system. The retrieved bacteria sample shows higher concentration of target bacteria than the original bacteria sample that has a lower concentration.

**Fig. 2.**
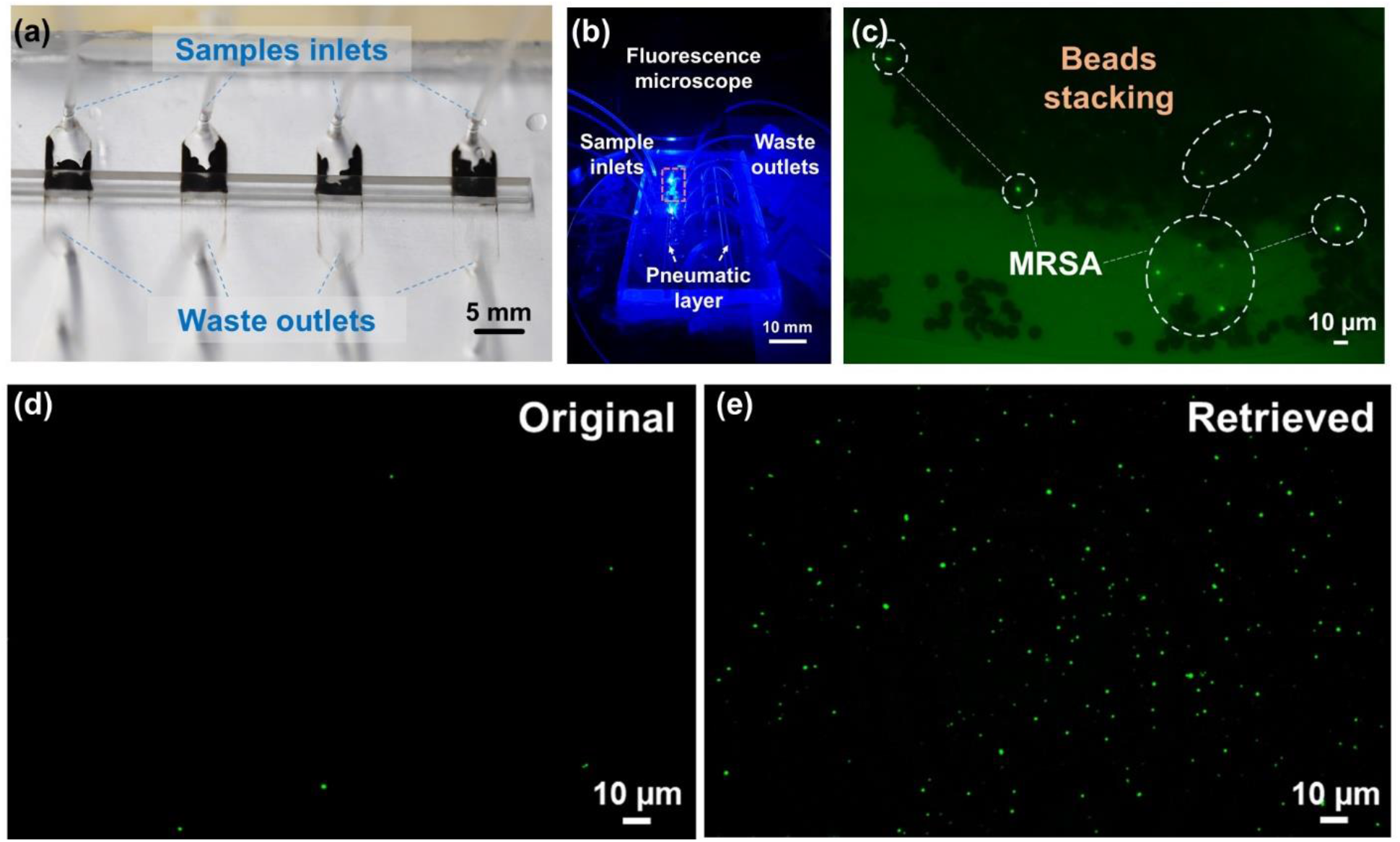
(a) Beads pattern in the channel without MRSA. (b) Experimental setup of multiplexing separation of target bacteria under the fluorescence microscope. (c) Beads stacking with MRSA under fluorescence microscope. (d) The original bacterial sample and (e) retrieved bacterial sample were compared to indicate the on-chip concentration capability of the nano-sieve device.

After successfully concentrating MRSA using nano-sieve, standard plate count was employed to quantify the concentration ability of nano-sieve. **Table 1** displays the concentration factors achieved with various inlet concentrations using the nano-sieve device. The concentration factor was determined by dividing the inlet concentration by the outlet concentration. A total of 600 μL of MRSA was injected into inlets, while 30 μL of PBS was used to retrieve the MRSA, resulting in a theoretical concentration factor of 20. However, as the MRSA concentration increased, the experimental concentration factor slightly decreased. One possible reason for this is that some bacteria may have leaked out through the waste outlets when the MRSA concentration was too high. This suggests that the nano-sieve is more suitable for concentrating low-concentration bacteria, aligning with our objective of enhancing the detection limit.

**Table 1.** Concentration factor of nano-sieve.

| Channel | Inlet Concentration (CFU/mL) | Outlet Concentration (CFU/mL) | Experimental Concentration Factor | Theoretical Concentration Factor |
| --- | --- | --- | --- | --- |
| 1 | $4.00 \times 10^2$ | $(6.00 \pm 2.94) \times 10^3$ | 15.00 | 20.00 |
| 2 | $5.93 \times 10^3$ | $(8.47 \pm 1.07) \times 10^4$ | 14.27 | 20.00 |
| 3 | $7.43 \times 10^4$ | $(8.37 \pm 1.40) \times 10^5$ | 11.26 | 20.00 |
| 4 | $3.43 \times 10^5$ | $(4.07 \pm 0.60) \times 10^6$ | 11.84 | 20.00 |
| 5 | $3.80 \times 10^6$ | $(4.07 \pm 0.60) \times 10^7$ | 10.79 | 20.00 |

**Fig. 3a** shows the nucleic acid purification process using a magnet after the MRSA lysis. In this process, magnet beads are introduced into a solution containing DNA, wherein a substantial amount of salt and polyethylene glycol (PEG) is present ^38^. The DNA molecules become crowded out and bind to the surface of the beads through electrostatic interactions ^39^ and molecular crowding ^40^. Magnet fields are then applied to collect DNA-bound beads, effectively removing unwanted debris such as membrane lipids and proteins ^41^. Then, guanidinium chloride is utilized to wash DNA as it disrupts protein-DNA interactions and aids in solubilizing and denaturing proteins ^42^. Finally, purified DNA is eluted using nuclease-free water. **Fig. 3b** shows the mechanism of RPA amplification. This process relies on the coordinated activities of recombinases, single-stranded DNA-binding proteins, and DNA polymerases to achieve isothermal amplification of the target DNA. The reaction is initiated by recombinases facilitating the binding of primers to the target DNA sequence. Single-stranded DNA-binding proteins stabilize the displaced DNA strands, allowing DNA polymerases to efficiently extend the primers along the DNA template in the presence of deoxynucleotide triphosphates (dNTPs), resulting in the synthesis of new DNA strands. Following this, the RPA amplicons are introduced into the CRISPRCas12a reaction. As shown in **Fig. 3c**, in positive samples, when the Cas12a-crRNA complex encounters the complementary target DNA, it undergoes a conformational change, leading to the activation of its nuclease activity. Cas12a then cleaves the target DNA at a specific site (cis-cleavage) as well as the collateral ssDNA probes (trans-cleavage), leading to the release of fluorescence signals from the fluorophore. In negative samples lacking the target DNA, the nuclease activity of Cas12a remains inactivated, preventing the cleavage of the probe and the generation of fluorescence signals. The assay development started with primer screening. Two CRISPR RNA and four primers sets were chosen, and their sequences are listed in **Table 2**. The input DNA was extracted from 10^8^ CFU/mL MRSA and purified using magnet beads. The excitation and emission wavelengths were set at 480 nm and 520 nm, respectively. The results of the primer screening are depicted in **Fig. 4a**. With the preceding RPA amplification, the fluorescence signal exhibited a substantial increase in comparison to CRISPR-Cas12a detection performed without RPA. Among the groups, the combination of crRNA2 and primer set 4 demonstrated the highest fluorescence signal, thus being chosen for subsequent experiments. To further evaluate the assay performance, a comparison was made between the magnet beads purification and a scenario without such purification. The results revealed a significant reduction in fluorescence signal without magnet beads purification, as illustrated in **Fig. 4b**. One possible explanation is that the presence of EDTA, lysosome, and proteinase K, which were introduced during MRSA lysis, might have disrupted the enzyme system, ultimately leading to the failure of DNA amplification and detection. On the other hand, magnetic beads purification effectively eliminated unwanted debris, including lysosome and proteinase K, thereby ensuring the successful amplification and detection of the target DNA. **Fig. 4c** displays the TEM images of magnet beads only (top) and magnet beads plus DNA (bottom). In the top image, aggregation and clustering of the beads can be observed, while the bottom image demonstrates the binding of DNA molecules to the magnet beads through electrostatic interactions and molecular crowding. Furthermore, the specificity performance of the assay was evaluated using three additional strains: wild-type *E. coli* K12 (**Fig. 4d**), kanamycin-resistant *E. coli* K12 (**Fig. 4e**), and wild-type S. aureus (**Fig. 4f**). The input DNA was extracted from bacterial cultures with a concentration of 10^8^ CFU/mL. The PBS buffer solution lacking bacteria was served as no template control (NTC). The results show that the fluorescence signals generated by these bacteria were comparable to that of the NTC group. However, upon mixing these bacteria with MRSA at a ratio of 1:1, the fluorescence signal of the mixture significantly increased. Particularly, the fluorescence signal produced by wild-type S. aureus reached nearly the same level as that in pure MRSA. These results not only confirm the specificity of the assay but also suggest that the presence of other bacteria does not interfere with MRSA detection results.

**Table 2.**
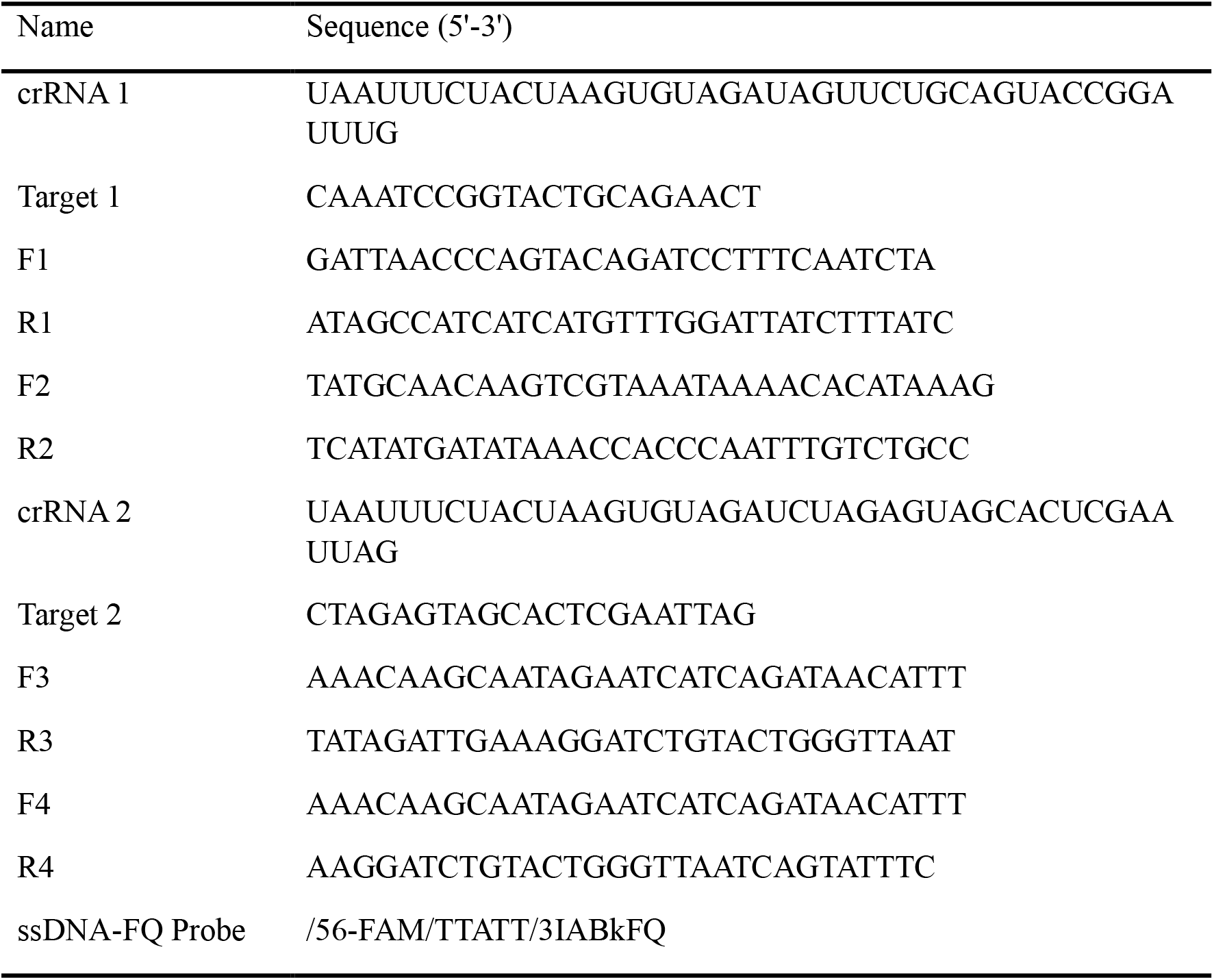
List of synthetic oligos sequence used in this study.

**Fig. 3.**
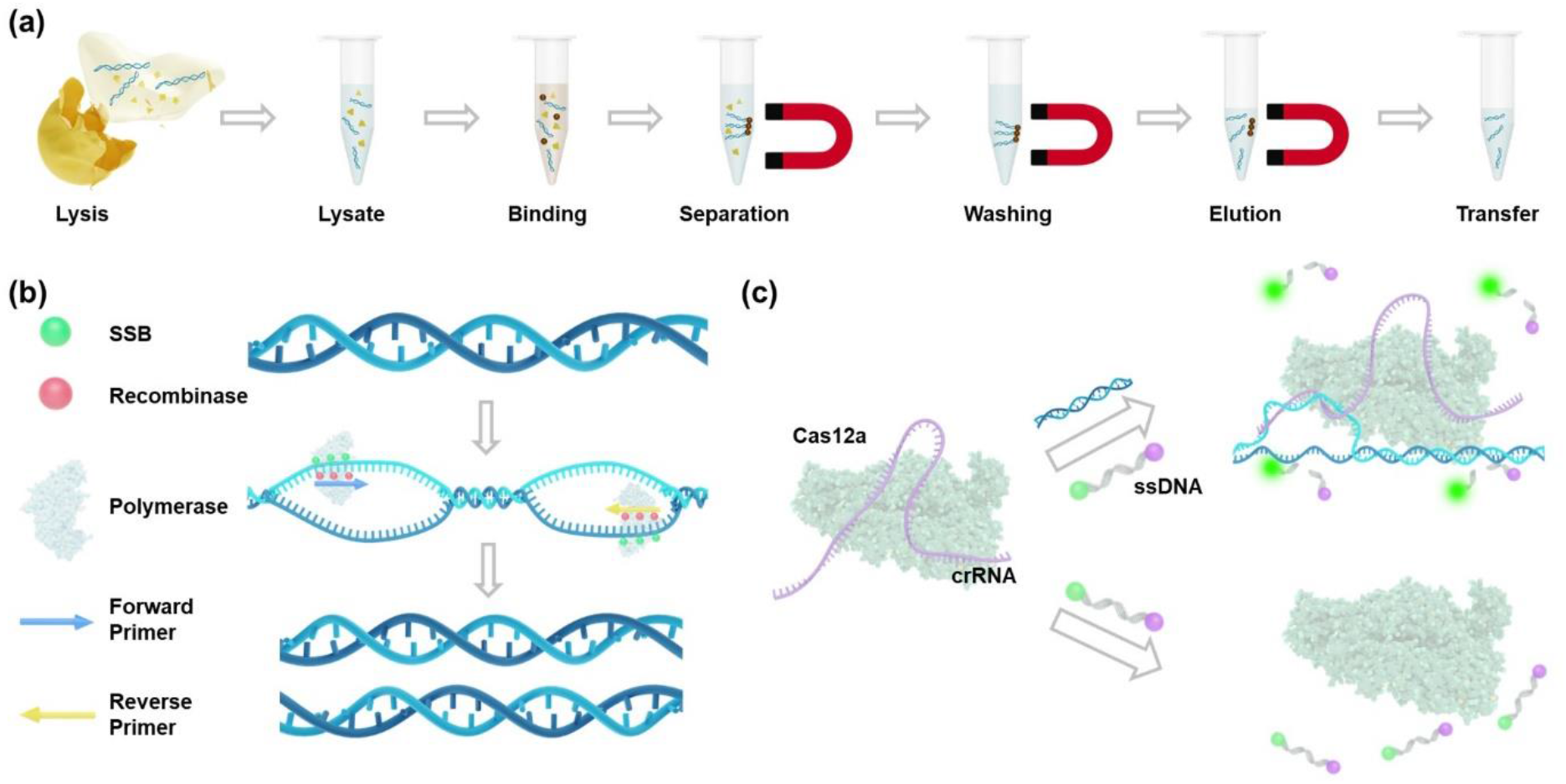
(a) Nucleic acid purification process using magnet beads. (b) Mechanism of RPA amplification. (c) Mechanism of CRISPR-Cas12a cleavage.

**Fig. 4.**
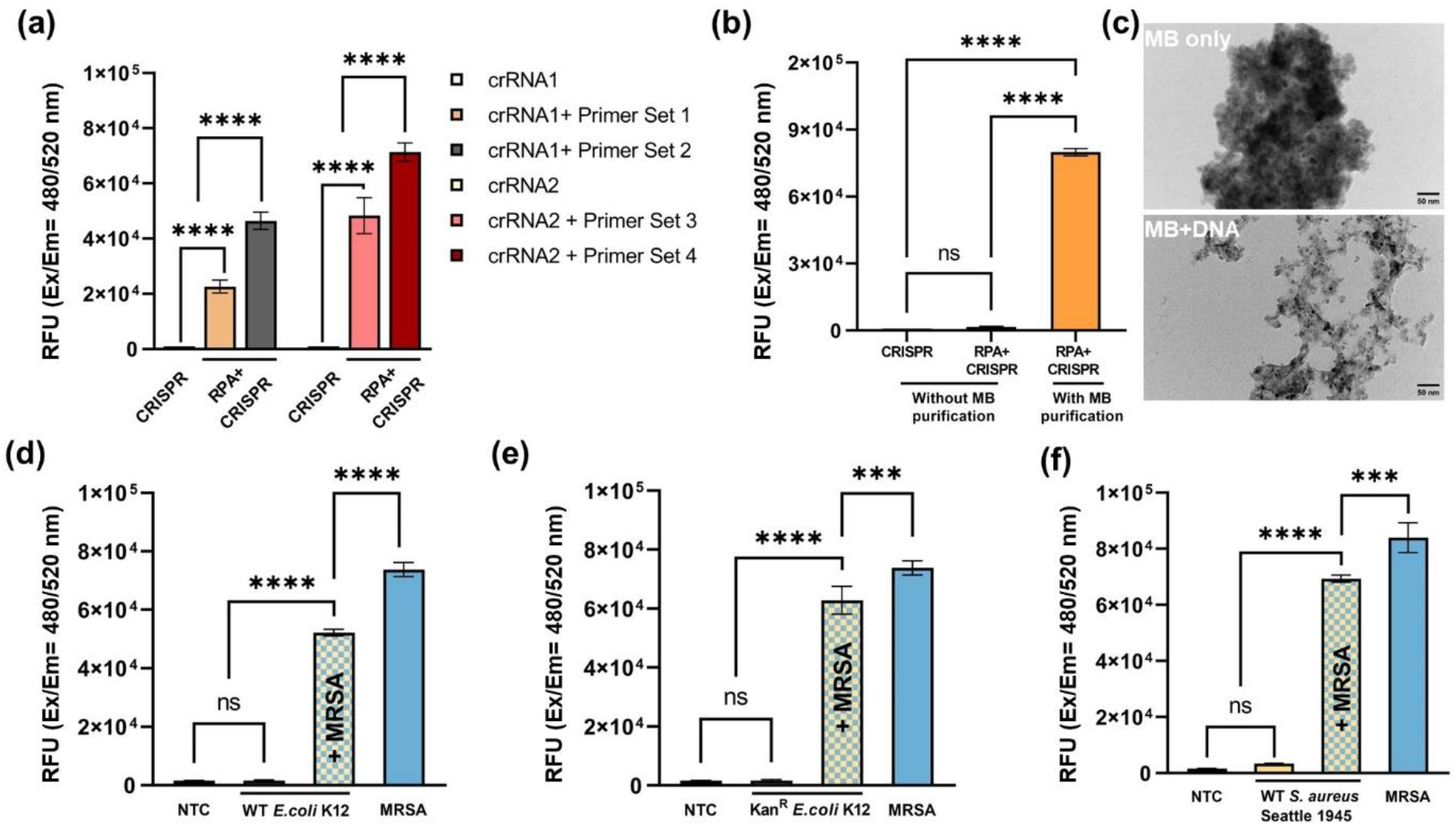
(a) Primer screening for RPA amplification. (b) MRSA detection with/without purification using magnet beads. (c) TEM images of magnet beads only (top) and magnet beads plus DNA (bottom), with a scale bar of 100 nm. (d), (e), and (f) represent the specificity tests of this assay using wild-type *E. coli* K12, kanamycin-resistant *E. coli* K12, and wild-type *S. aureus*, respectively. “NTC” refers to no template control. The data are represented as mean ± standard deviation (n = 3). For statistical analysis, ns, not significant = p > 0.05; *** = 0.0001 < p ≤ 0.001; **** = p ≤ 0.0001.

**Fig. 5a** presents a comparison of fluorescence signal obtained from off-chip and on-chip detection, encompassing various inlet MRSA concentrations ranging from 10^4^ CFU/mL to 10^6^ CFU/mL. When the MRSA concentration reached 10^6^ CFU/mL, both the on-chip and off-chip results exhibited saturation in fluorescence signal. As the MRSA concentrations decreased, the off-chip results displayed a decline in fluorescence signal, whereas the on-chip results remained saturated. Notably, at an MRSA concentration of 10^4^ CFU/mL, a discernible difference in fluorescence signal between the on-chip and off-chip results became evident, as depicted in **Fig. 5b**. Subsequently, the on-chip detection limit was determined by further decreasing the MRSA concentration. **Fig. 5c** presents the fluorescence signal acquired from on-chip detection using inlet MRSA concentrations ranging from 10^1^ CFU/mL to 10^3^ CFU/mL, accompanied by the corresponding endpoint images displayed in **Fig. 5d**. At an MRSA concentration of 100 CFU/mL, the naked eye easily discerned the fluorescence differences between the positive and negative groups, which were further validated through one-way ANOVA analysis of quantified characterization ^43^. Therefore, a detection limit of 100 CFU/mL was established for MRSA detection using the nano-sieve device.

**Fig. 5.**
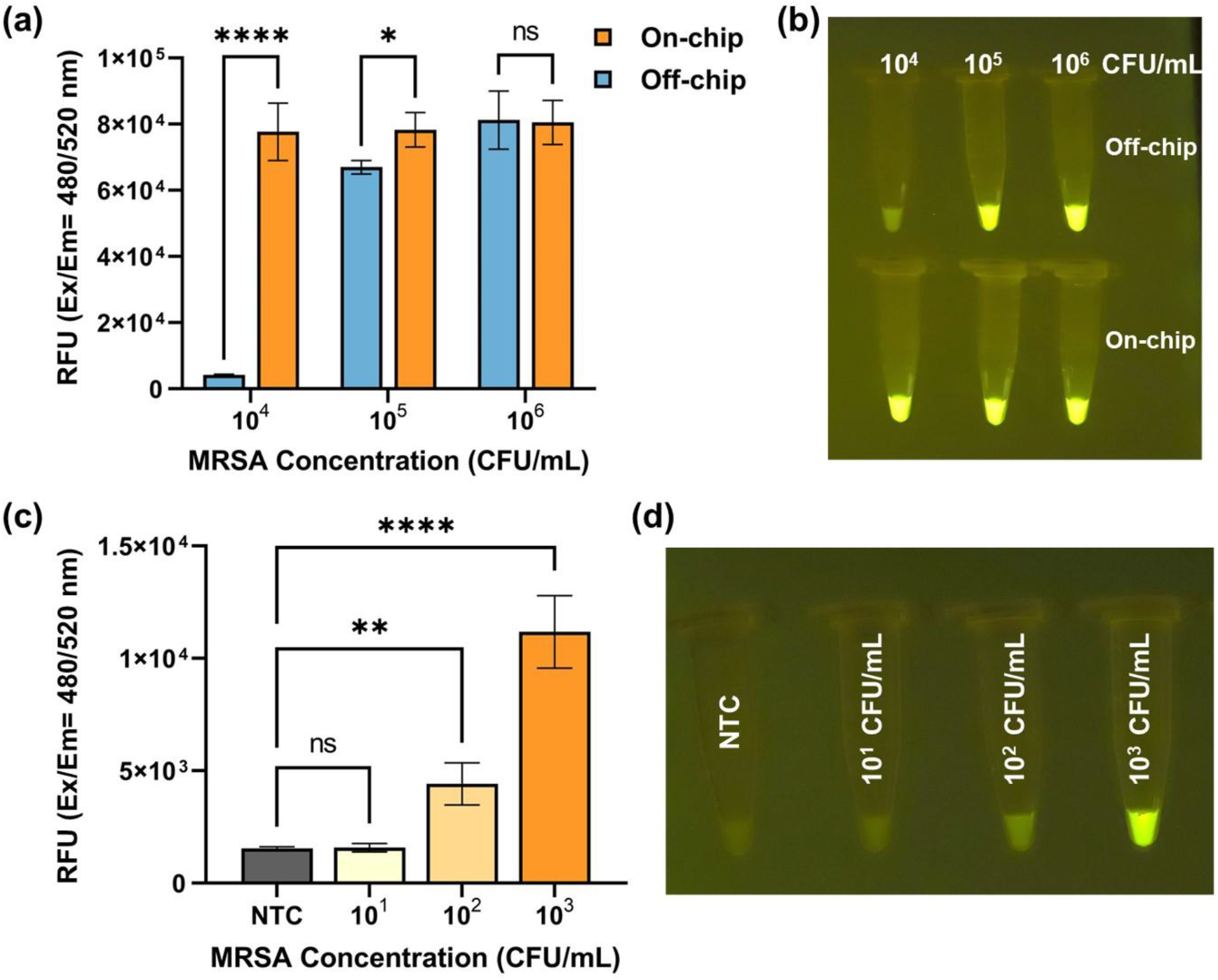
(a) Quantified fluorescence signal of off-chip and on-chip detection with varying inlet MRSA concentrations ranging from 10^4^ CFU/mL to 10^6^ CFU/mL (Ex/Em = 480/520 nm). (b) Endpoint images of the reactions excited by a transilluminator (wavelength: 465 nm) in response to different inlet MRSA concentrations ranging from 10^4^ CFU/mL to 10^6^ CFU/mL (excitation wavelength: 465 nm). (c) Quantified fluorescence signal of on-chip detection with inlet MRSA concentration ranging from 10^1^ CFU/mL to 10^3^ CFU/mL (Ex/Em = 480/520 nm). (d) Endpoint images of the reactions excited by a transilluminator (wavelength: 465 nm) in response to different inlet MRSA concentrations ranging from 10^1^ CFU/mL to 10^3^ CFU/mL. “NTC” refers to no template control. The data are represented as mean □} standard deviation (n = 3). For statistical analysis, ns, not significant = p > 0.05; *** = 0.0001 < p ≤ 0.001; **** = p ≤ 0.0001.

The approach of rapidly purifying and highly concentrating pathogens from a large volume of bodily fluids could be crucial for disease diagnostics, such as sepsis, at the early stage ^44,45^. Compared to the surface chemistry technique, pathogen separation based on the physical structure of microfluidic platforms could be simpler and more efficient, while minimizing contamination issues. Our nano-sieve system has been functionalized via a 3-D beads-stacked microstructure that can be precisely tuned by applied flow rate ^46^, and optimized via a pneumatic layer that can counterbalance the hydrodynamic pressure during the flow condition ^47^. Moreover, our pneumatically-regulated nano-sieve has been developed with an extremely low aspect ratio of 1:25,000, significantly reducing the hydrodynamic pressure that may affect the mechanically-driven separation process. This flexible pneumatic layer enhances the adaptability of our nano-sieve channel compared to a rigid nanofluidic channel. Therefore, this deformable pneumatic layer enables a more reliable bead-stacking by applying the positive air pressure during the purification process and an easier target release by offering the negative air pressure during the retrieval process. In addition, we recently showed that the pneumatic-controlled nano-sieve with a patterned microstructure on the bottom of the substrate could efficiently enhance the capture of nanoscale targets at a higher flow rate ^47^. In the future, the development of 3-D spaced beads array, such as a combination of various sized beads, could be beneficial to a higher capture efficiency of target bacteria, aiming to improve the detection limit by combining with our optimized molecular detection technique.

While our current study involved spiking MRSA into plasma samples, future research will focus on testing clinical samples to validate the applicability of our approach in a real-world clinical setting. Currently, our group is engaged in the development of a rapid and efficient method for purifying and identifying antibiotic-resistant bacteria (ARB) from human blood samples. Through an immunomagnetic assay, highly concentrated red blood cells (RBCs) could be removed from bacteria spiked samples, while effectively retaining the target bacteria for subsequent purification using the optimized nano-sieve device. Presently, our system only processes up to six samples simultaneously, but by introducing more channels during the chip fabrication process, we can achieve the simultaneous detection of hundreds of clinical samples within a 6-inch wafer using a simple equipment setup comprising a multipump, pipetting system, microcentrifuge, heat block, magnet, transilluminator, and necessary reagents. Such high-throughput capability will significantly reduce the turnaround time for patients and healthcare providers to obtain the test results, which is crucial for expediting disease diagnosis, facilitating prompt medical interventions, and ensuring timely implementation of appropriate treatment strategies ^48^. Additionally, by designing different primers and CRISPR RNA, this microfluidic device enables multiplexing for different pathogens. This feature holds significant promise in the diagnosis of disease potentially caused by multiple pathogens, such as sepsis ^49^. While our detection assay is currently performed in tubes after the nano-sieve concentration, requiring manual pipetting, our future work will focus on the incorporation of a platform for the detection assay, such as the funnel-adapted sensing tube (FAST) ^50^ or a digital multiplex dRPA chip ^51^. Also, we could simplify the operation process by introducing a one-pot RPA and CRISPR assay ^52^.

## Conclusion

In conclusion, we have introduced a miniaturized and versatile nano-sieve device with a pneumatically-regulated chamber for the rapid purification and highly concentrated isolation of MRSA from plasma samples. Our simplified and cost-efficient fabrication process, incorporating multiple channels and a 3D beads-stacked microstructure, enables highly efficient capture of MRSA by precisely controlling the flow rate, resulting in a significant concentration of captured bacteria. The integration of this device with the RPA and CRISPR-Cas12 assay enhances the detection sensitivity, achieving a lower on-chip detection limit of 100 CFU/mL compared to the off-chip limit of 10^4^ CFU/mL. Our sensitive detection method can be completed within a short timeframe of 4 hours under physiological temperature conditions, eliminating the need for centrifugation. The scalability of the nano-sieve device allows for the simultaneous processing of multiple clinical samples and multiplexing detection of different pathogens. By improving the sensitivity and specificity of MRSA detection, our approach holds promise in contributing to better patient outcomes and addressing the challenges posed by antibiotic-resistant bacterial infections.

## Experimental Methods

### Device fabrication

The pneumatically-regulated nano-sieve device consists of a channel-patterned glass substrate and a PDMS topping with a pneumatic chamber. Briefly, a cleaned 4-inch glass wafer (University wafer, D263, 550 μm, double side polished) was deposited by a layer of TEOS (PECVD, AME P5000) with a thickness of 200 nm. After spin-coating a thin layer (1 μm in thickness) of positive photoresist (AZ Mir 701), a plastic photomask (Fineline Imaging, CO, USA) was applied to transfer the channel pattern onto the photoresist layer. Followed by buffer oxide etching (BOE) procedure, the nano-sieve channel was created on the glass substrate. To fabricate a pneumatic chamber of 2 mm in height and 2 mm in width, a PDMS mixture with one part of curing agent and ten parts of base polymer (SYLGARDTM 184, Krayden Inc., CO, USA) was poured into a three-dimensional (3-D) printed mold (Fictiv, CA, USA). After the curing process of PDMS in the oven at 60 °C overnight, the chamber layer was punched by a 1 mm puncher (INTEGRATM MiltexTM), for the pneumatic regulation by the air pump (Precigenome LLC, CA, USA). This chamber layer was bonded onto a thin layer of cured PDMS thin film (200 μm in thickness) via plasma treatment (Electro-Technic Products, IL, USA). Then, the entire part was baked on the hot plate at 100 °C for 2 hrs to obtain the robust bonding in between. To finalize the fabrication of a functional device, the nano-sieve channel on the substrate was sealed by bonding with the pneumatic layer via plasma treatment and baking process on a hot plate. Before using the device for experiments, the holes of inlet and outlet were punched by a 1 mm puncher, so that the microfluidic tubing (Scientific Commodities,Inc., BB31695-PE/3) can connect the device to the sample sources in a syringe (BD 1 mL, NJ, USA).

### Beads stacking functionalization

A strategy of creating the 3-D microstructure of stacked beads was applied to physically capture the target bacteria from the initial sample. In this study, the nano-sieve channel was first rinsed and cleaned by isopropyl alcohol (IPA) solution. Before introducing the magnetic beads into the channel, the air pump was used for ensuring the pressure applied in the pneumatic layer consistently at 12 Psi. Then, a 50 μL of 10 μm magnetic beads with a concentration of 10 mg/mL was injected into the channel to form a “coarse filter” by employing a syringe pump with a flow rate of 30 μL/min. The injection of another 50 μL of 5 μm magnetic beads with a concentration of 10 mg/mL into the channel was followed, to form a “fine filter” under a flow rate of 20 μL/min. Finally, this functionalized nano-sieve device is ready for separating the target bacteria from the introduced sample solution.

### Bacterial culture

Methicillin-resistant *Staphylococcus aureus* (ATCC 43300) and *Staphylococcus aureus* (ATCC 25923) were purchased from Fisher Scientific, Kanamycin-resistant *E. coli* 10798 and *E. coli* MG1655 were from lab stock. The bacteria were cultured in tryptic soy broth medium (MilliporeSigma) and maintained on the tryptic soy agar plate. After the overnight culture at 37 °C under 200 rpm, 1 mL of bacterial cells was centrifuged at 8,000g for 5 min to form a pellet. The pellet was then resuspended in 1 mL of phosphate buffer saline (PBS) or a mixture of 1:4 of human plasma (MilliporeSigma) and PBS. This process was repeated twice to completely remove culture media. The concentration of the bacterial cells was determined by counting the colony-forming unit (CFU) on standard TSA plates. The harvested cells were diluted in PBS or a 1:4 mixture of plasma and PBS using a 10-fold dilution for future use.

### Bacterial staining

Two microliters of BacLight green dye (MilliporeSigma) was mixed with 1 mL PBS-based bacteria solution, and the mixture was incubated at room temperature for 20 min by following the manufacturer’s instruction. Then, the solution was centrifuged at 8,000g for 5 min to discard the PBS that consists of extra dye. The bacteria were resuspended in 1 mL fresh PBS for the further use.

### Bacteria capture and concentration

The 600 μL of prepared sample solutions, in which the bacteria with a certain concentration were spiked in 1:4 diluted plasma solution, was first loaded into several 1 mm sterile syringes. A multi-channel syringe pump was used to simultaneously introduce the sample solutions into the functionalized nano-sieve channels through the microfluidic tubing, under a flow rate of 4 μL/min. After completing the separation process, the air pump was stopped and a 30 μL of fresh PBS solution was applied for rinsing the entire channel to retrieve the magnetic beads and the captured targets, from the channel to a sterile centrifuge tube. A simple and direct separation with an external magnet was used for extracting the bacteria involved PBS solution, which is ready for the further detection based on RPA/CRISPR technique. The concentration factor in this case is expected to be 20-fold.

### Fluorescence microscopy imaging

After the completion of bacteria separation, 10 μL of each retrieved sample and initial sample were placed onto a glass slide for the measurement of cell density. Under the microscope equipped with a high-speed camera and the fluorescent light resource, the fluorescent images were captured and analyzed by using Leica LAS X software.

### TEM Characterization

A JEOL 2100 transmission electron microscope (TEM) operating at 200 kV was used to image the samples. An aqueous solution of magnetic beads +/- DNA was deposited onto a carbon-coated copper TEM grid and left to allow evaporation of the water. Then, the dried grids were dipped into 18 Mega Ohm water for 30 s to remove excess salts. The images were then captured with a Gatan Orius camera.

### Nucleic acid extraction

The nucleic acid preparation involves bacterial lysis and DNA purification, which was carried out at room temperature and completed in less than 30 min. To prepare the enzymatic lysis buffer (20 mM Tris-HCl, 2 mM EDTA, and 1.2% Triton X-100), lysozyme (Thermo Fisher) was added at a concentration of 20 mg/mL immediately before use. Next, 18 μL of the enzymatic lysis buffer was added to 10 μL of the pathogens and incubated for 10 min. Following this, 2 μL of proteinase K (Thermo Fisher) was added, and the mixture was incubated for 5 min. For DNA purification, AMPure XP beads (Beckman Coulter) were used following the vendor’s instruction with slight modification. Briefly, 54 μL of magnet beads were added to the lysate and incubated for 5 min. The tube was then placed on the magnet for 2 min to separate beads from the solution. Next, the beads were washed twice with 70 μL of 5 M Guanidinium chloride (MilliporeSigma), and the DNA was eluted with 20 μL of nuclease-free water (Thermo Fisher).

### RPA amplification and CRISPR-Cas12a detection

TwistAmp^@^ Basic kit was purchased from TwistDx™. The RPA primers, crRNA, AsCas12a, and fluorophore-quencher probes were all obtained from Integrated DNA Technologies, and detailed information about the synthetic oligonucleotides are listed in **Table 2**. The RPA primer sets were designed using PrimerQuest™ Tool. Additionally, NEBuffer™ r2.1 was purchased from New England Biolabs. The RPA reaction was conducted based on the instructions: A mixture of 29.5 μL of rehydration buffer, 11.2 μL of nuclease-free water, and 2.4 μL each of forward and reverse primers (10 μM) was added to the enzyme pellet. Then, 2 μL of purified DNA and 2.5 μL of MgOAc (280 mM) were added and mixed to achieve a total volume of 50 μL. The mixture was incubated at 37 °C for 20 min. Following the incubation, 2 μL of RPA amplicons were added to a pre-assembled CRISPR-Cas12a mixture comprising 50 nM of AsCas12a, 62.5 nM of crRNA, 10× buffer, and 2.5 μM of ssDNA-FQ probe, resulting in a final reaction volume of 20 μL. The reaction solution was incubated at 37 °C for 30 min. After the incubation, the mixture was excited by a blue light transilluminator (brand: SmartBlue, Part number: NEB-E4100, excitation wavelength of 465 nm) for naked-eye observation. Finally, 20 μL of nuclease-free water was added to 5 μL of the mixture, which was then characterized by an Agilent BioTek Cytation 5 imaging reader.

## Author Contributions

Ruonan Peng, Xinye Chen and Ke Du designed the experiments. Ruonan Peng, Xinye Chen, Fengjun Xu, and Richard Hailstone performed the experiments. Ruonan Peng and Xinye Chen wrote the manuscript. All authors commented on the manuscript.

## Acknowledgements

This study was supported by The National Institutes of Health R35GM 142763.

